# *Akkermansia Muciniphila* induces chronic extramedullary hematopoiesis through cooperative IL-1R and TLR signals

**DOI:** 10.1101/2022.01.03.474846

**Authors:** Yuxin Wang, Tatsuya Morishima, Maiko Sezaki, Gaku Nakato, Shinji Fukuda, Yuhua Li, Hitoshi Takizawa

## Abstract

Bacterial infections can activate and mobilize hematopoietic stem and progenitor cells (HSPCs) from the bone marrow (BM) to spleen, which is termed as extramedullary hematopoiesis (EMH). Recent studies suggest that commensal bacteria, particularly the microbiota, regulates not only the host immune system but also hematopoietic homeostasis. However, the impact of gut microbial species on hematopoietic pathology remains largely unknown. Here we found that systemic injection of *Akkermansia muciniphila* (*A. m*.), a mucin-degrading bacterium abundantly existing in the human gut rapidly activates BM myelopoiesis, and induces a slow but long-lasting hepato-splenomegaly, characterized by the expansion and differentiation of functional HSPCs, which we termed chronic EMH. Genetic deletion of Toll-like receptor-2 and -4 (TLR2/4) partially diminished *A. m*.-induced chronic EMH, while additional pharmacological inhibition of the interleukin-1 receptor (IL-1R) completely alleviated splenomegaly and EMH. Our results demonstrate that cooperative IL-1R- and TLR-mediated innate immune signals regulate commensal bacteria-driven EMH, which might be relevant for certain autoimmune disorders.

**Article Summary:** The aim of our study is to understand how *Akkermansia muciniphila (A*.*m*.), one of the major mucin-degrading microbial species in the human gut activate the immune and hematopoietic systems in a mouse model. We found that a single injection of the *A*.*m*. membrane fraction can induce long-lasting hepatosplenomegaly with splenic EMH through cooperative IL-1R- and TLR-mediated innate immune signals.

## Introduction

Commensal microbes, which coexist within the human body and reach a cell number more than total human cells^1^, ca. 4×10^13^, have been recently shown to significantly affect the maintenance and activation of the immune system^2, 3^. The microbiota colonized in the gut has also been shown to regulate hematopoietic development^4^, steady-state hematopoiesis^5, 6^, and hematopoietic stem cell (HSC) aging^7^. *Akkermansia muciniphila* (*A. m*.) is an anaerobic gram-negative bacterium and one of the major mucin-associated microbial species found in the human gut, which accounts for 1-5% of the entire microbial community^8^. They reside in the mucus layer as a mucin-degrading bacterium^9^, and their increase can alleviate metabolic syndromes^10^, inflammatory bowel diseases (IBD)^11^ and PD-1-based immunotherapy against epithelial cancer^12^. In contrast to these beneficial roles, *A. m*. has been associated with intestinal inflammation^13^ and increased GVHD-related mortality^14^, suggesting additional pathological roles in immune system control. However, little is known regarding its direct effect on the hematopoietic system and its pathogenesis.

In steady state, adult HSCs are homeostatically maintained in the bone marrow (BM), where specific niche cells support HSC maintenance and quiescence, while only a very minor population of HSCs can be found in other organs such as the peripheral blood (PB)^15^, lymph duct^16^, spleen^17^ and gingiva^18^. In contrast, under hematopoietic stress conditions such as infection^19^ or chemotherapy^20^, HSCs are activated and mobilized from the BM to other organs such as the spleen and liver to enhance hematopoiesis and replenish cell types that are in high demand, which is called extramedullary hematopoiesis (EMH). EMH is known to be mediated by cytokine/chemokine receptors that sense the inflammatory milieu released from proximal/distal organs, including granulocyte colony-stimulating factor receptor (G-CSFR)^19^, c-Kit, CXCR4^20^, and pattern recognition receptors (PRRs) such as Toll-like receptors (TLRs), NOD-like receptors (NLRs)^19^ and C-type lectin receptors (CLRs)^21^.

In addition to PRRs, another innate immune sensor is the interleukin (IL)-1 receptor, which binds to IL-1α and IL-1β and triggers downstream transcriptional responses via the adaptor protein MyD88^22^. While IL-1β is secreted systemically and circulates throughout different tissues, IL-1α is produced locally in inflamed tissues by various hematopoietic (e.g. monocytes^23^, granulocytes, and inflammatory Ly-6C^high^ cells^7^) and non-hematopoietic cells. IL-1α is crucial for sustaining inflammatory responses^24^, recruiting myeloid cells to infected tissue^25^ and inducing hematopoietic stem and progenitor cell (HSPC) mobilization and expansion both *in vitro* and *in vivo*^26, 27^. Yet, the contribution of IL-1 to EMH is so far unknown.

In this study, we found that 1) single administration of the *A. m*. membrane fraction induces drastic, long-lasting anemia and hepato-splenomegaly with immune cell and HSPC activation, 2) there are two temporal waves of HSPC expansion causing EMH upon *A. m*. injection, the latter with functional HSPCs accumulated in the spleen, 3) *A. m*.-induced EMH is mediated entirely by MYD88/TRIF-dependent innate immune signals, and partially by TLR2/4 dependent signals, and 4) pharmacological inhibition of IL-1R in addition to TLR2/4 genetic ablation completely alleviates EMH. These novel findings might be involved in the pathophysiology of *A. m*.-associated metabolic and inflammatory disorders, and IL-1-related chronic hemato-immune disorders such as stem cell aging, clonal hematopoiesis and autoimmune diseases.

## Methods

### Mice

C57BL/6, B6.SJL, *Hlf*^*tdTomato+*^ reporter^28^, *Myd88;Trif*^-/-^ and *TLR2;4*^-/-^ mice were maintained at the Center for Animal Resources and Development at Kumamoto University. Experiments were approved by the Animal Care and Use Committee of Kumamoto University.

### *A. muciniphila* preparation

*A. muciniphila* was cultured in GAM medium (Nissui Pharmaceutical) under anaerobic conditions. After 2 to 3 days of culture, bacteria were collected, sonicated using a sonicator (Smurt, Microtec) and ultracentrifuged at 100,000 x g for 60 minutes at 4°C (Optima XE-90, Beckman). Protein concentration of the bacterial membrane suspension was determined with Bio-Rad protein assay (BioRad).

### Treatment

200μg of LPS, 200μg of Pam3CSK4, 200μg of *A. m*. or PBS were intraperitoneally (*i*.*p*.) injected into mice. For the IL-1R inhibition experiment, mice were *i*.*p*. injected daily with 37μg of Anakinra (Kineret, IL-1R antagonist, Swedish Orphan Biovitrum AB) over 2 weeks until sacrificed for analysis. *A. m*. was injected once, one day after the first Anakinra injection.

### FACS analysis

BM and spleen were harvested from treated mice and incubated with biotinylated antibodies against the lineage (Lin) markers: B220, CD3*ε*, CD4, CD8*εα*, NK1.1, CD11b, Ter119, and Gr-1 together with the fluorescence-conjugated antibodies: c-Kit, Sca-1, CD34, Flt3, CD150, CD48, CD16/32, IL-7R*εα* and CD86; for mature cell analysis, with: B220, CD3*ε*, F4/80, Ly6G, Ly6C, CD11c and CD11b. Stained cells were analyzed on FACS Canto II or FACS Aria III flow cytometers (BD Biosciences). Data was analyzed using FlowJo (BD Biosciences).

### Competitive repopulation assay

3×10^5^ whole BM (WBM) cells and 1×10^6^ whole splenocytes were isolated from either *A. m*.-or PBS-treated WT mice (CD45.1^+^) at 14 days post injection and transplanted intravenously into lethally-irradiated (10Gy), 8-10 weeks old WT mice (CD45.2^+^) together with 3×10^5^ WBM competitor cells isolated from WT mice (CD45.1/CD45.2). PB donor chimerism was assessed every 4 weeks, up to 16 weeks and in BM up to 20 weeks post transplantation.

### Measurement of inflammation-or HSC niche-related factors

BM and spleen fluid were diluted with BSA/PBS (0.1%). Splenocyte lysate was treated in 500µM 1x RIPA buffer (Abcam). G-CSF was quantified by Mouse G-CSF ELISA Kit (Proteintech) according to the manufacturer’s instructions. Other cytokines were measured by Legendplex™ Mouse Inflammation Panel (BioLegend) and Legendplex™ Mouse HSC Panel (BioLegend) according to the manufacturer’s instructions.

### Quantification and statistical analysis

Statistical analysis was performed using SPSS version 22. Two-group comparisons were analyzed with the Student’s t-Test (two-tailed t-test). Multigroup comparisons were performed by a one-way analysis of variance (ANOVA) followed by the Tukey–Kramer multiple comparisons test if variances were equal, and Dunnett’s C test if variances were not equal. All results are expressed as means ± standard error of the mean (SEM). The criterion of significance was set at *p* < 0.05.

## Results

### Single injection of *A. m*. induced long-lasting anemia and hepatosplenomegaly with extramedullary hematopoiesis

To study the impact of *A. m*. on the hematopoietic system, we *i*.*p*. injected the *A. m*. membrane fraction into wild-type (WT) mice and analyzed different hematopoietic organs. PB counts showed pancytopenia with sudden decrease in white blood cell (WBC) and platelet (PLT) counts at day 1 post injection (Fig. 1A). Intriguingly, *A. m*.-treated mice showed sustained severe anemia and hepato-splenomegaly, which peaked at day 14 and lasted for almost 2 months after a single injection (Fig. 1B-C). The total cell number of splenocytes followed the same kinetics (Fig. 1D).

**Figure 1.**
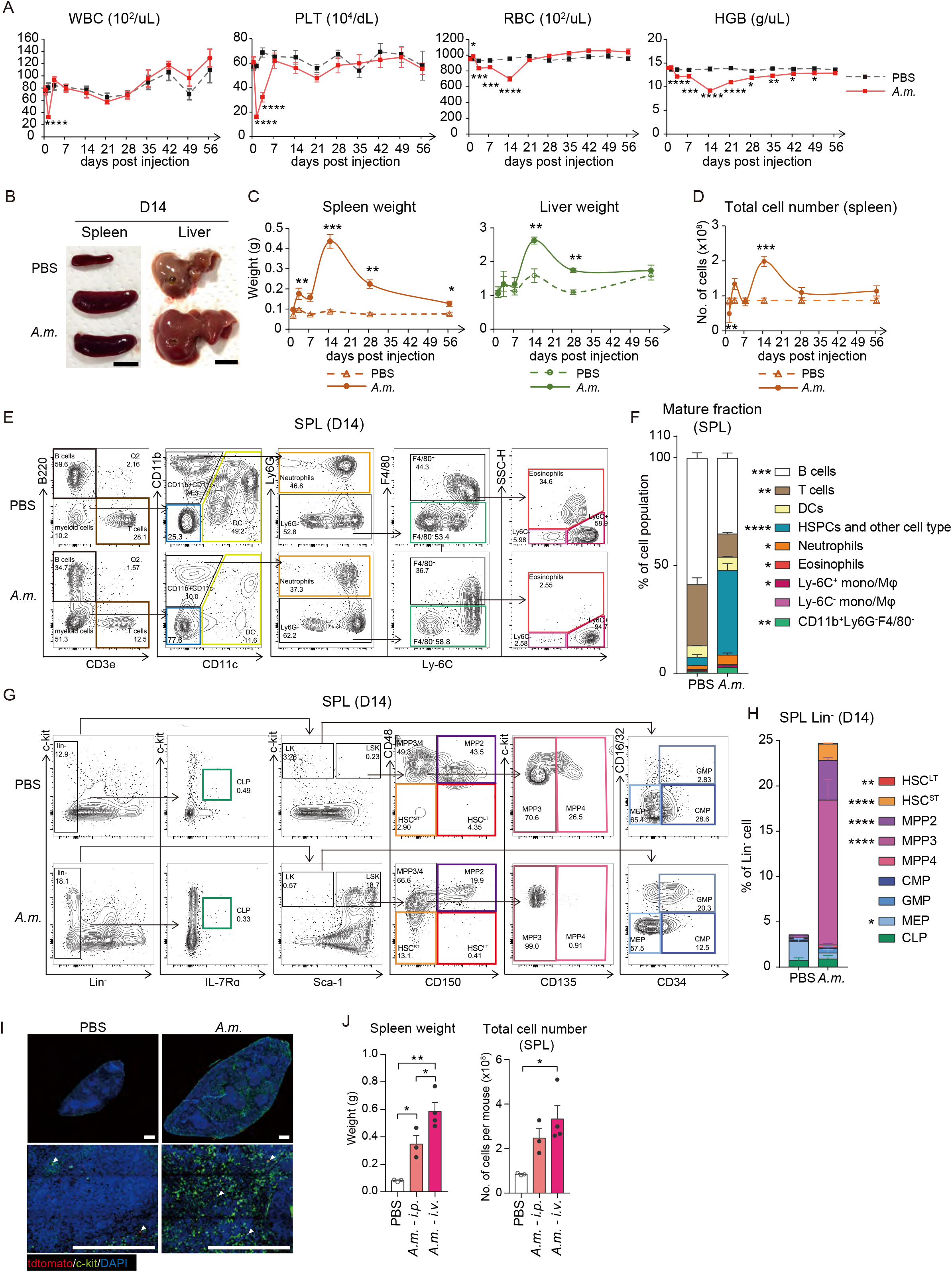
A single shot of *A. m*. injection induced long-lasting anemia and hepatosplenomegaly with extramedullary hematopoiesis. (A) Time-course kinetics of peripheral blood counts of PBS-or *A. muciniphila* (*A. m*.)-injected mice. WBC, white blood cell; PLT, platelet; RBC, red blood cell; HGB, hemoglobin (n=5-20 from 3-8 independent experiments). (B) Representative images of the spleen and liver at day 14 after *A. m*. injection. Scale bars indicate 1cm. (C) Time-course kinetics of spleen and liver weights after PBS or *A. m*. injection (n=4-8 from 2-5 independent experiments). (D) Total nucleated cell number of splenocytes after PBS or *A. m*. injection (n=4-16 from 2-10 independent experiments). (E-F) Representative FACS plots (E) and frequency (F) of mature hematopoietic cells in the spleen (SPL) 14 days after PBS or *A. m*. injection (n=12-13 from 7 independent experiments). (G-H) Representative FACS plots (G) and frequency (H) of HSPC subpopulation (LK, LSK and CLP) in hematopoietic lineage negative cells in spleen at 14 days after PBS or *A. m*. injection (n=5-8 from 4 independent experiments). (I) Representative images of the spleen from PBS (left)-or *A. m*. (right)-treated *Hlf*^*tdTomato+*^mice at day 14 (c-Kit, green; tdTomato, red; DAPI, blue). Arrowheads indicate putative HSCs (c-Kit^+^tdTomato^+^). Scale bars indicate 500µm. (J) Spleen weight and total cell number of splenocytese at day 14 after *A. m*. injection via *i*.*p*. versus *i*.*v*. routes (n=3 from 2 independent experiments). \**p*<0.05, \*\**p*<0.01, \*\*\**p*<0.001, \*\*\*\**p*<0.0001 (two-tailed t-test).

To dissect which cells expanded in *A. m*.-treated spleen, mature and immature hematopoietic cell fractions were immunophenotyped at day 14 post injection (Fig. 1E and S1A). Myeloid cells such as neutrophils (CD11b^+^Ly6G^+^), eosinophils (CD11b^+^Ly6G^-^F4/80^+^SSC-H^high^) and Ly-6C^+^ monocytes (CD11b^+^Ly6G^-^F4/80^+^SSC-H^low^Ly-6C^+^) expanded significantly in *A. m*.-injected spleen, whereas B and T cells decreased compared to` PBS controls (Fig. 1E-F). The population with the most significant increase was CD3*ε*^-^B220^-^CD11b^-^CD11^-^ (blue gate), containing both non-hematopoietic cells and immature HSPCs, with a fold expansion of 8.5 (4.72±1.71% in control *vs* 39.97±4.31% in *A. m*.-injected group). This was not observed in the BM (22.40±3.23% in control *vs* 25.50±3.70% in *A. m*.-injected group) (Fig. S1B-C).

The spleen is a site for EMH in response to systemic infection. Past studies report HSPC mobilization to the spleen upon injection of live *E*.*coli* or its membrane components, as well as upon infiltration of the microbiota induced by colitis^19, 29, 30^. Based on previous reports and our above-mentioned observations, we hypothesized that long term-sustained splenomegaly is driven by the mobilization and expansion of HSPCs in the spleen. When splenocytes were stained with the respective HSPC makers, the frequency of hematopoietic lineage negative (Lin^-^) cells increased significantly in *A. m*.-injected mouse compared controls at day 14 post injection (*p*<0.001, 3.66±0.44 % in control *vs* 24.80±2.51% in *A. m*.-injected group) (Fig. 1G-H). In contrast, there was little change in the Lin^-^ fraction of *A. m*.-treated BM (31.30±1.38% in control *vs* 31.83±2.12% in *A. m*.-injected group) (Fig. S1D-E). Within the Lin^-^ fraction, Lin^-^Sca-1^+^c-Kit^+^ (LSK) cells containing hematopoietic stem cells (HSCs) and multipotent progenitors (MPPs), both of which have self-renewal and multi-lineage differentiation potential, significantly expanded in *A. m*.-injected BM and spleen (Fig. 1G-H and S1D-E). Supporting this data, immunofluorescent imaging of the spleen of *Hlf*^*tdTomato*^ mice, that allow to visualize putative HSPC^28^ showed expansion of c-Kit^+^tdTomato^+^ HSPCs and Gr-1^+^/Ly6G^+^ mature myeloid cells in the red pulp of *A. m*.-injected spleen compared with controls (Fig. 1I and S1F-G).

Given *A. m*. infiltrates through the gut mucosa and reaches the BM through the circulation^7^, we compared different injection routes of *A. m*. treatment. Intravenous (*i*.*v*.) injection of *A. m*. induced a slightly bigger splenomegaly (Fig. 1J) and comparable HSPC expansion both in the spleen and BM (Fig. S1H), suggesting that the splenomegaly was induced by *A. m*. irrespective of the injection route.

Collectively, these data indicate that a single shot of *A. m*. induced significant hematopoietic responses characterized by long term-sustained anemia and hepato-splenomegaly with EMH.

### Time-course kinetic analysis revealed two differential HSPC waves in *A. m*.-injected spleen

To characterize the cellular basis for long-term, sustained splenomegaly upon *A. m*. injection, time-course kinetics of HSPC populations were next analyzed. The LSK fraction showed rapid increase in the BM (2.14±0.15% in control *vs* 9.88±1.13% in *A. m*.-injected group) as well as PB (0.22±0.03% in control *vs* 1.27±0.21% in *A. m*.-injected group) one day after *A. m*. injection, suggesting HSPC mobilization (Fig. 2A-B). This “first wave” of HSPC-containing LSK increase in the BM and PB appeared to cease within a week as shown by the frequency of LSK cells returning back to baseline by day 7 (Fig. 2A-B). Subsequently, the Lin^-^Sca-1^-^c-Kit^+^ (LK) fraction containing myeloid lineage-committed progenitors without self-renewal potential^31^ gradually increased in the spleen (4.92±1.66% in control *vs* 18.64±1.72% in *A. m*.-injected group) and reached its peak at day 7, followed by their differentiation into mature myeloid cells (Ly6C^+^ monocyte, dendritic cells (DCs), neutrophils) peaking at day 14 (Fig. 2D and S2A). After the “first wave”, a delayed but stronger “second wave” of HSPC increase (called chronic EMH thereafter) was observed with a peak at day 14 in *A. m*.*-*injected spleen (0.56±0.14% in control *vs* 23.08±2.27% in *A. m*.-injected group) and BM (2.14±0.15% in control *vs* 18.90±2.18% in *A. m*.-injected group) (Fig. 2A-B). Subpopulation analysis of the delayed chronic phase showed none of the LSK subfractions maintained their increase in the spleen nor BM, but gradually returned to baseline by day 56 (Fig. 2D and S2B-C).

**Figure 2.**
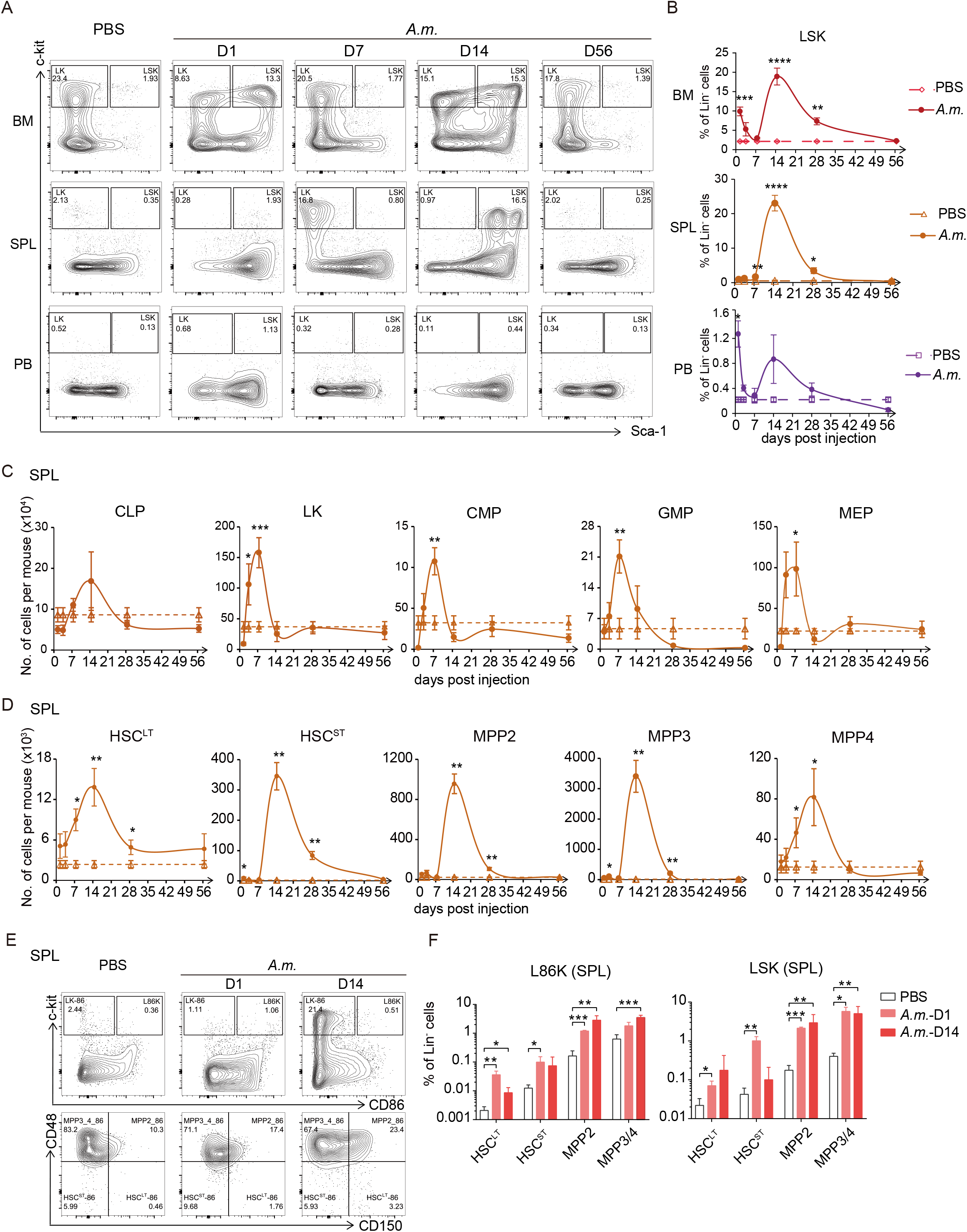
Time-course kinetic analysis revealed two distinct waves of HSPC increase in the spleen of *A. m*.-injected mice. (A-B) Representative FACS plots of Lin^-^ cells (A) and percentage of LSK cells (B) in the BM (upper), spleen (middle), and PB (lower) *A. m*.at indicated timepoints after PBS or *A. m*. injection. (C) Absolute number of CLP and LK subfractions in the spleen of PBS or *A. m*.-injected mice. (D) Absolute number of HSPC subfractions within LSK in the spleen of PBS or *A. m*. injected mice (3-14 mice from 3-8 independent experiments). (E-F) Representative FACS plots (E) and frequency (F) of CD86-or Sca-1-defined HSPCs in the spleen at day 1 and 14 post PBS or *A. m*. injection (n=3-8 from 5 independent experiments). \**p*<0.05, \*\**p*<0.01, \*\*\**p*<0.001, \*\*\*\**p*<0.0001 (two-tailed t-test).

The HSPC marker Sca-1^32^ is one of the interferon (IFN)-responsive genes known to be upregulated upon bacterial infection and thus, may result in a spillover of Sca-1 negative LK into LSK cells and thereby overestimation of HSPC number^33^. CD86 has been recently reported as a surrogate HSPC marker under infection-stressed conditions^34^. When we compared the frequency of L86K (Lin^-^ CD86^+^c-Kit^+^)-defined HSPCs with that of LSK-defined HSPCs at day 1 and day 14 after *A. m*. injection (Fig. 2E-F and S2D-E), both HSPC identification methods resulted in an overall comparable numbers in the BM and spleen after *A. m*. stimulation.

Collectively, the following data indicates two distinct waves of HSPC increase, a quick-responsive “first wave” and a delayed but long-lasting “second wave” in *A. m*.*-*injected spleen.

### *A. m*.*-*expanded HSCs in the spleen are transplantable

Functional HSCs are detectable in the spleen in steady-state and under stress conditions^17, 19^. To examine whether HSCs that accumulated in *A. m*.-stimulated spleen are functional, we performed a competitive transplantation assay (Fig. 3A). At 14 days after *i*.*p*. treatment with either PBS or *A. m*. in WT mice (CD45.1^+^), 3×10^5^ WBM and 1×10^6^ splenocytes were transplanted into lethally-irradiated WT recipient mice (CD45.2^+^), together with 3×10^5^ WBM competitors from untreated WBM (CD45.1^+^CD45.2^+^). Monthly donor chimerism of the PB revealed that *A. m*.*-*treated WBM donors showed multilineage reconstitution comparable to that of PBS-treated controls over 4 months post transplantation (Fig. 3B). However, *A. m*.*-*treated splenocytes showed significantly higher engraftment levels in total hematopoietic cells (CD45^+^) and myeloid lineage cells (CD11b^+^Gr-1^+^) but not in B or T cells (Fig. 3C). This was further confirmed by terminal BM analysis at 5 months post transplantation, with a significantly higher donor chimerism in HSCs and MPPs of the BM transplanted with *A. m*.*-* treated splenocytes compared to controls (Fig. 3D). In contrast, no significant differences could be observed in WBM donor transplants. These data indicate that consistent with immunophenotypic analysis (Fig. 2), functionally transplantable HSCs possibly with myeloid-biased lineage output increased in the spleen at day 14, after one shot of systemic *A. m*. injection.

**Figure 3.**
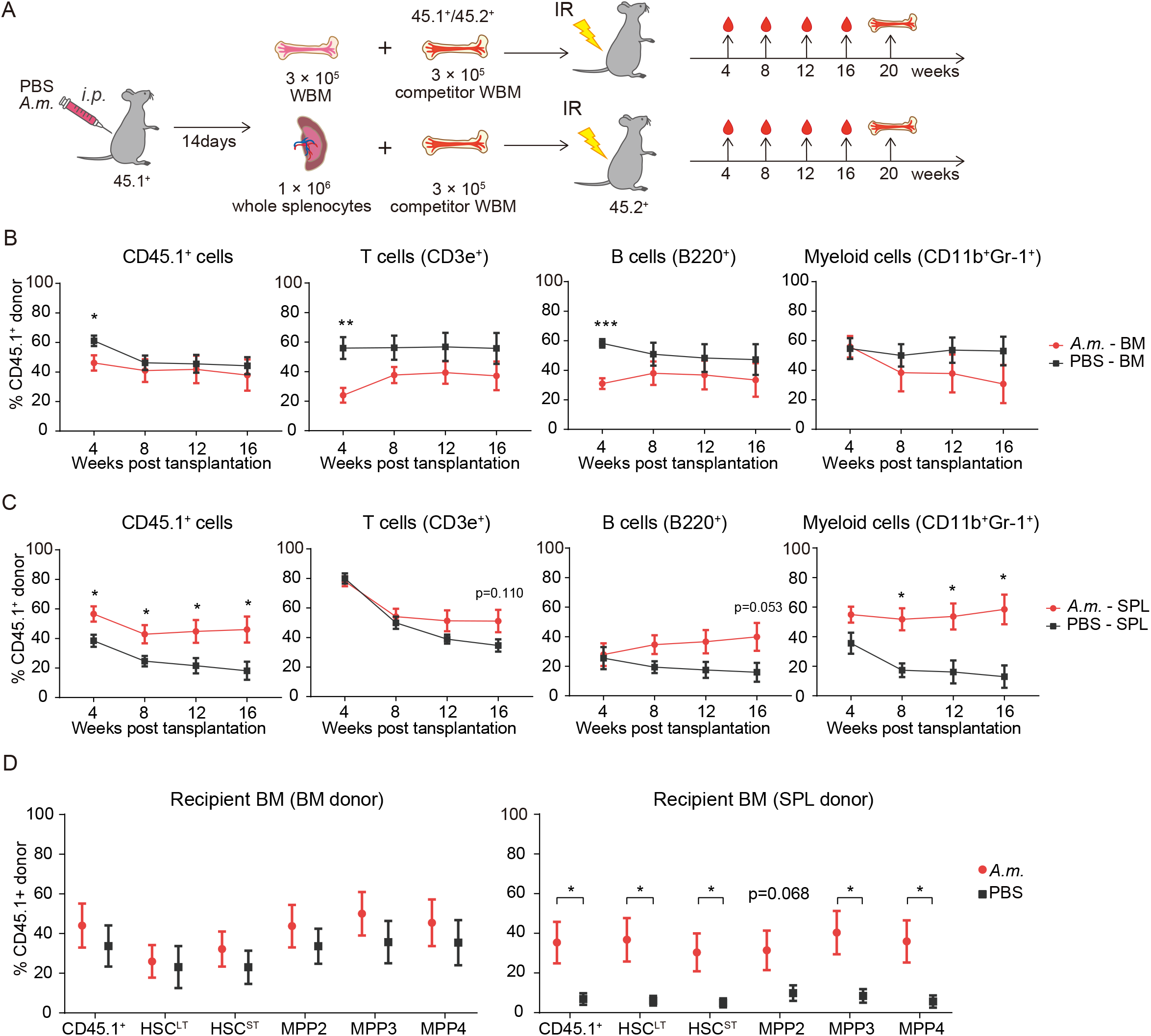
*A. m*. stimulation expanded functional HSCs in the spleen. (A) Experimental scheme of competitive transplantation: 3×10^5^ whole bone marrow (WBM) cells or 1×10^6^ whole splenocytes were isolated from PBS or *A. m*.*-*treated WT mice (CD45.1^+^) at day 14 post injection, mixed with 3×10^5^ WBM competitors (CD45.1/CD45.2^+^) and transplanted into lethally-irradiated (10Gy) WT mice (CD45.2^+^). Donor chimerism in the PB was analyzed monthly for up to 16 weeks and the BM for 20 weeks. (B-C) Time course kinetics of donor chimerism in the PB of recipients transplanted with whole BM (B) or splenocytes (C). Shown are donor chimerisms of leukocytes (CD45.1^+^), myeloid (CD11b^+^Gr-1^+^), B-lineage (B220^+^), and T-lineage (CD3*ε*^+^) cells. (D) Donor chimerism in HSPCs from the BM of recipients transplanted with whole BM (left) or splenocytes (right) (n=7-11 from 2 independent experiments). \**p*<0.05, \*\**p*<0.01, \*\*\**p*<0.001 (two-tailed t-test).

### *A. m*.-induced HSPC accumulation in the spleen is partially mediated by TLR2/4 and entirely by MYD88/TRIF dependent signals

*A*.*m*. is a gram-negative bacterium with a membrane that contains lipopolysaccharide (LPS)^10^, which has the potential to activate and mobilize HSCs directly through the Toll-like receptor (TLR) 4-signaling pathway^19^. The purified membrane protein, Amuc_1100 isolated from the *A*.*m*. membrane fraction has been reported to bind to TLR2^10^. Therefore, we next tested whether *A. m*. is able to recruit HSPCs in the spleen via TLRs. *Tlr2;4*^-/-^ double knock out (DKO) mice lacking TLR2 and TLR4 expression and *Myd88;Trif* ^-/-^ DKO mice lacking the down-stream adaptor molecules of TLR and IL-1R signaling were challenged with PBS or *A. m*. and analyzed 14 days later (Fig. 4A). The severe splenomegaly that occurred in WT mice was partially and completely cancelled in *Tlr2;4*^-/-^ DKO and *Myd88;Trif* ^-/-^ DKO, respectively (Fig. 4B-C), with little to no HSPC accumulation in the spleen and BM (Fig. 4D-E and S3A-B). These results indicate that *A. m*.*-*induced EMH is partially dependent on TLR2/4-mediated signals but completely dependent on MYD88/TRIF-mediated signals.

**Figure 4.**
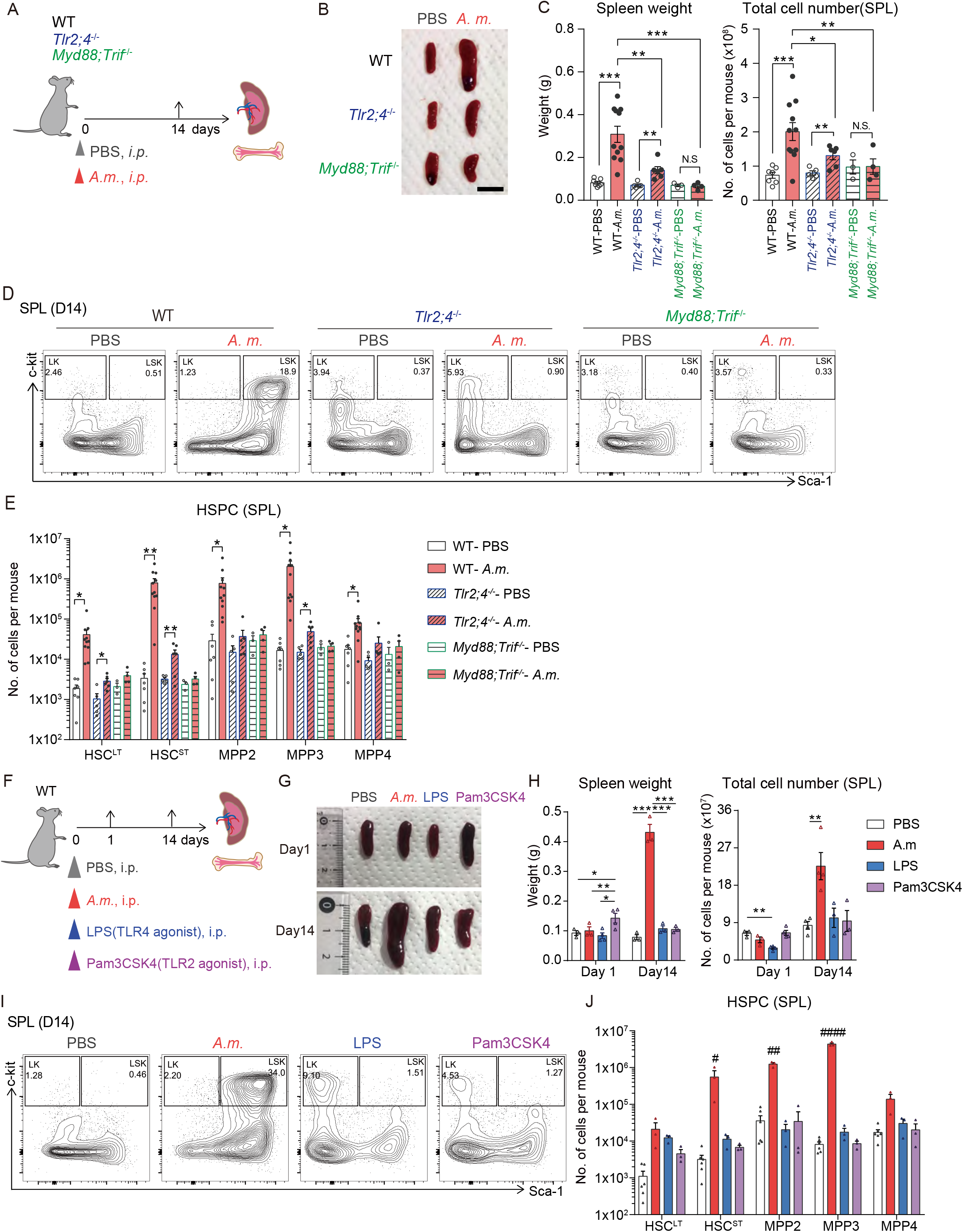
*A. m*.*-*induced HSPC expansion in the spleen is mediated by innate immune signals, but not entirely by TLR2/4 signals. (A) Experimental scheme of *A. m*. injection into WT versus innate immune signal-deficient mice: WT, *Tlr2*;*4* or *Myd88;Trif* DKO mice were *i*.*p*. treated with PBS or *A. m*. and the spleen and BM were analyzed at day 14 post injection. (B) Representative images of the spleen. Scale bar indicates 1cm. (C) Spleen weight (left) and total cell number (right) of splenocytes from WT mice treated with PBS (WT-PBS) or *A*.*m*. (WT-*A*.*m*.), *Tlr2;4* DKO mice injected with PBS (*Tlr2;4* DKO-PBS) or *A*.*m*. (*Tlr2;4* DKO-*A*.*m*.), or *Myd88;Trif* DKO mice injected with PBS (*Myd88;Trif* DKO-PBS) or *A*.*m*. (*Myd88;Trif* DKO-*A*.*m*.) at day 14 post injection (n=4-11 from 4 independent experiments)WT, *Tlr2*;4 or *Myd88;Trif* DKO mice treated with PBS or *A. m*. (D) Representative FACS plots of Lin^-^ splenocytes from PBS or *A. m*.*-*treated WT, *Tlr*2;4 or *Myd88;Trif* DKO mice at day 14 post injection. (E) Absolute number of HSPC subpopulations in the spleen of WT mice injected with PBS (WT-PBS) or *A. m*. (WT-*A. m*.), *Tlr2;4* DKO mice injected with PBS (*Tlr2;4* DKO-PBS) or *A. m*. (*Tlr2;4* DKO-*A. m*.), or *Myd88;Trif* DKO mice injected with PBS (*Myd88;Trif* DKO-PBS) or *A. m*. (*Myd88;Trif* DKO-*A. m*.) at day 14 post injection (n=4-11 from 4 independent experiments). (F) Experimental scheme of *A. m*. versus TLR agonist treatment: WT mice were *i*.*p*. injected with PBS, 200μg *A. m*., 200μg LPS or 200μg Pam3CSK4, and the spleen and BM were analyzed 14 days later. (G) Representative images of the spleen treated with PBS, *A. m*., LPS, or Pam3CSK4 at day 1 (upper) and 14 (lower) post injection. (I) Spleen weight (left) and total cell number of splenocytes (right) at day 1 and 14 (n=3-7 from 6 independent experiments). (I-J) Representative FACS plots (H) of Lin^-^ cells and absolute cell number (J) of HSPC subpopulations of the spleen at day 14. (n=4-7 from 2 independent experiments). \**p*<0.05, \*\**p*<0.01, \*\*\**p*<0.001 (two-tailed t-test). ^#^*p*<0.05, ^##^*p*<0.01, ^###^*p*<0.001, ^####^*p*<0.0001 (one-way ANOVA). N.S., not significant.

To further confirm splenomegaly-inducing signals, LPS and Pam3CSK4, which are TLR4 and TLR2 agonists, respectively were *i*.*p*. injected into WT mice and compared to effects caused by the same dosage of *A. m*. (Fig. 4F). Although a single shot of Pam3CSK4 induced mild splenomegaly at day 1, neither LPS nor Pam3CSK4 could induce splenomegaly and HSPC accumulation in the spleen and BM comparable to *A. m*. at day 14 (Fig. 4G-J and S3C-D). Additionally, the combination of LPS and Pam3CSK4 also failed to trigger splenomegaly (data not shown).

Taken together, these data suggest that although TLR2/4 signals partially contribute to splenomegaly, additional TLR2/4 independent innate immune signals are required for triggering *A. m*.-induced splenomegaly.

### Sustained elevation of splenic IL-1α production contributes to TLR2/4-independent chronic EMH

We found that MYD88/TRIF is fully responsible, while TLR2/4 is partially for *A. m*.-induced EMH (Fig. 4C). Since other TLRs are also involved in MYD88/TRIF-mediated pathways^35, 36^ and proinflammatory cytokines, such as IL-1^37^, interferons (IFNs)^32, 38^, tumor necrosis factor-α (TNF-α)^39^ affect HSC fate and blood cell output, we considered the possibility of EMH mediated through cytokines. Therefore, we next quantified various inflammatory factors in the serum, spleen fluid and spleen cell lysate isolated from *A. m*.-treated mice (Fig. 5A-C and S4A-D). Among all the cytokines measured, the concentration of IL-1α showed a remarkable 25-fold increase at day 1 and 2-fold increase at day 14 in the spleen fluid and spleen cell lysate of *A. m*.-treated mice compared to their corresponding controls (Fig. 5A and S4A). The elevation of IL-1α in the spleen was partially reduced in *Tlr2;4*^-/-^ DKO spleen and completely cancelled in *Myd88;Trif* ^-/-^ DKO spleen (Fig. 5A and S4A), suggesting partial TLR2/4- and complete MYD88/TRIF-dependent IL-1α secretion in the spleen. As IL-1α was not upregulated in the serum and likewise, undetectable in the BM (Fig. 5B and data not shown), its secretion appears restricted to the spleen. Although IL-1β and TNF-α also showed remarkable elevation in *A. m*.-treated spleen at day 1, their levels were not sustained for 2 weeks but appears totally dependent on both TLR2/4 and MYD88/TRIF signals (Fig. 5A and S4A). In contrast, IFN-*γ* showed a continuous increase over time in both the spleen and serum (Fig. 5A and S4A-B), while independent of TLR2/4- and MYD88/TRIF signaling. Measurement of G-CSF and CXCL12, both known as HSPC mobilization factors^19^ revealed transient upregulation of G-CSF in the serum and downregulation of CXCL12 in the BM fluid already at day 1 (Fig. 5C). These results indicate that both signals trigger initial HSPC mobilization during the early phase (Fig. 2A-B), but do not contribute to *A. m*.-induced chronic EMH.

**Figure 5.**
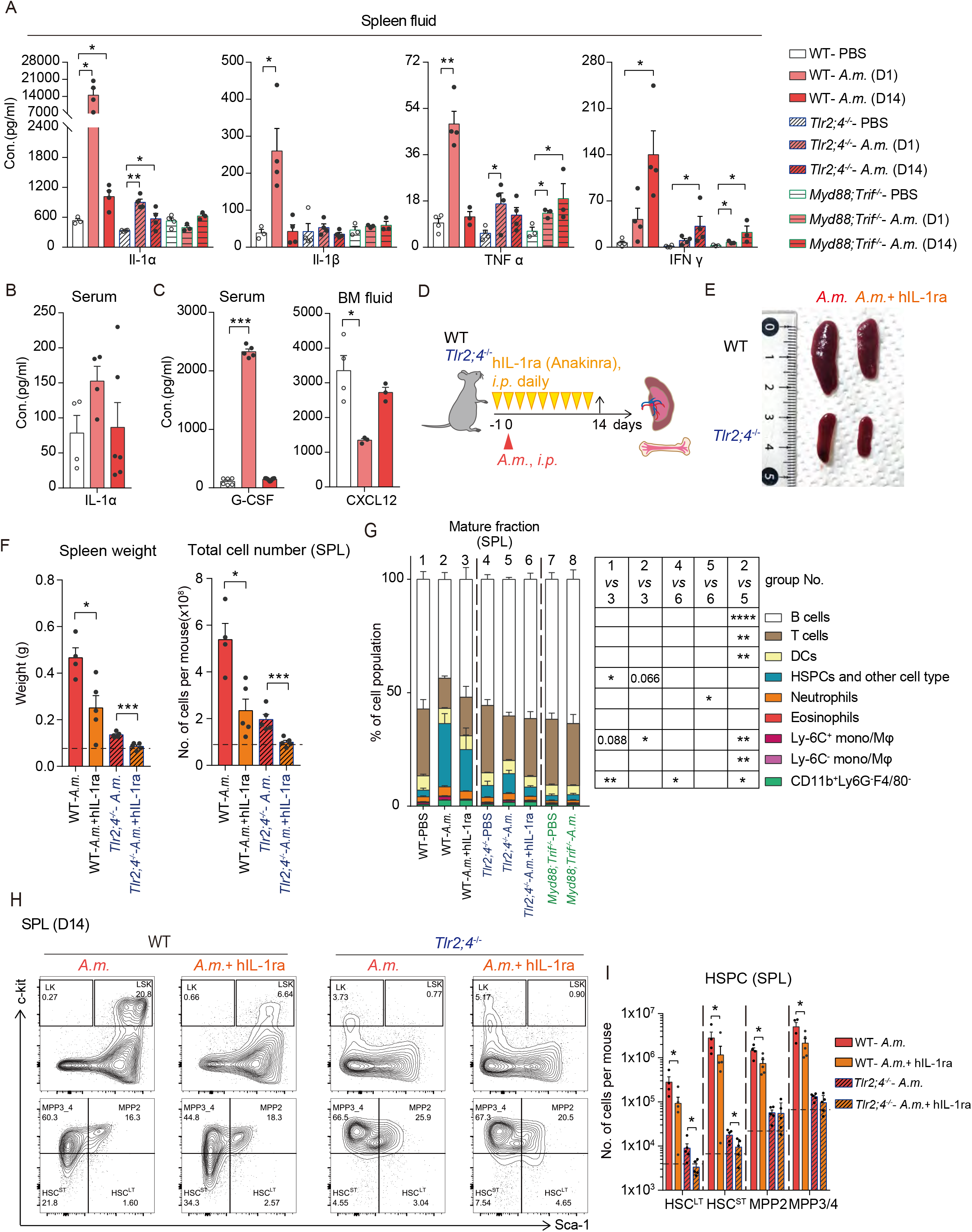
Local secretion of IL-1α contributes to chronic HSPC expansion in the spleen independent of TLR2/4 signaling pathways. (A) Concentration of IL-1α, IL-1*ε*, TNF-α and IFN-*γ* in spleen fluid from WT, *Tlr2;4* or *Myd88;Trif* DKO mice treated with PBS or *A. m*. at day 1 and 14 (n=3-4 from 4 independent experiments). (B-C) Concentration of IL-1α (B), G-CSF (C) in the serum and CXCL12 in the BM fluid (C) from WT mice treated with PBS or *A. m*. at day 1 and 14 (n=3-6 from 3 independent experiments). (D) Experimental scheme of hIL-1ra treatment: WT and *Tlr2;4* DKO mice were treated with *A. m*. followed by daily injections with or without hIL-1ra treatment. Spleen and BM were analyzed at day 14. (E) Representative images of the spleen from WT or *Tlr2;4* DKO treated with *A. m*. +/-hIL-1ra. (F) Spleen weight (left) and total cell number (right) of splenocytes at day 14 (n=5-6 from 4 independent experiments). Dashed line indicates spleen weight and cell numbers in PBS-treated spleen. (G) Frequency of mature hematopoietic cells in the spleen 14 days after PBS or *A. m*. injection. Statistical comparisons are shown in the box: *p* values that are between 0.05-0.1 are shown, blanks indicate *p*>0.1. (H-I) Representative FACS plots (H) and absolute cell number (I) of HSPC subpopulations in the spleen at day 14 (n=5-6 from 4 independent experiments). Dashed lines indicate absolute cell numbers of each HSPC fractions in PBS-treated spleen. \**p*<0.05, \*\**p*<0.01, \*\*\**p*<0.001, \*\*\*\**p*<0.0001 (two-tailed t-test).

Since IL-1α levels best correlated with splenomegaly and HSPC expansion that occurred in the spleen (Fig. 4C-E and 5A), it is likely that the remaining TLR2/4-independent hematopoietic responses are mediated by IL-1α signaling. To test this, we administered human IL-1R antagonist (hIL-1ra), Anakinra, daily to WT and *Tlr2;4*^-/-^ DKO mice followed by *A. m*. injection (Fig 5D). Anakinra treatment resulted in significant reduction of splenomegaly and the total cell number in both mouse strains, close to WT (ca. 59.0% reduction of spleen weight in WT-*A. m*.+hLI-1ra *vs* WT-*A. m*., while 86.6% reduction in *Tlr2;4*^-/-^-*A. m. vs* WT-*A. m*.) (Fig. 5E-F). Detailed analysis showed decrease of myeloid lineage cells and all LSK subpopulations in the spleen of both strains to near baseline level in *Tlr2;4*^-/-^ DKO mice (Fig. 5G-I). *A. m*.-induced splenomegaly and HSPC and myeloid cell accumulation in the spleen were partially cancelled in WT mouse and almost completely cancelled in *Tlr2;4*^-/-^ DKO mice (ca. 48.8% reduction of HSPC population in *Tlr2;4*^-/-^-*A. m*.+hLI-1ra *vs Tlr2;4*^-/-^-*A. m*.), indicating that cooperative IL-1R- and TLR2/4-mediated signals contribute to *A. m*.-induced chronic EMH. Of note, normalization of HSPCs by IL-1 inhibition was found only in the spleen but not in the BM (Fig. S4E-F), which again confirms local IL-1α signaling as key in the spleen.

Collectively, these data suggest that chronic local IL-1α production in the spleen upon *A. m*. treatment is regulated partly by TLR2/4, together with MYD88/TRIF signals, and cooperative interplay between TLR2/4 and IL-1R is required for local HSPC and myeloid cell expansion, ultimately leading to chronic EMH.

## Discussion

This study demonstrates that a single injection of *A. m*.-derived agents triggers a series of hematopoietic responses leading to chronic splenomegaly, sustained by two temporally-distinct waves of EMH (Fig. 6). At day 1 post *A. m*. injection, HSPCs rapidly increase in the BM and are mobilized to the PB via upregulation of G-CSF in the serum and downregulation of CXCL12 in the BM (Fig. 5C). The spleen showed pancytopenia and local production of IL-1α for emergency hematopoiesis (Fig. 2A-B). This “first wave” of HSPC increase appears to cease by their gradual differentiation into myeloid lineage-committed progenitors at day 7, followed by the generation of mature myeloid cells including Ly6C^+^ monocytes, DCs, and neutrophils by day 14 (Fig. 2D and S2A). After the “first wave” of EMH, a gradual but stronger “second wave” of HSPC expansion occurs at day 14 in the spleen and BM (Fig. 2A-B). Chronic accumulation of HSPCs and the expansion of mature myeloid cells result in sustained splenomegaly and long-term EMH. Moreover, we found that *A. m*. stimulation induced consistent local secretion of IL-1α in the spleen which induced chronic EMH.

**Figure 6.**
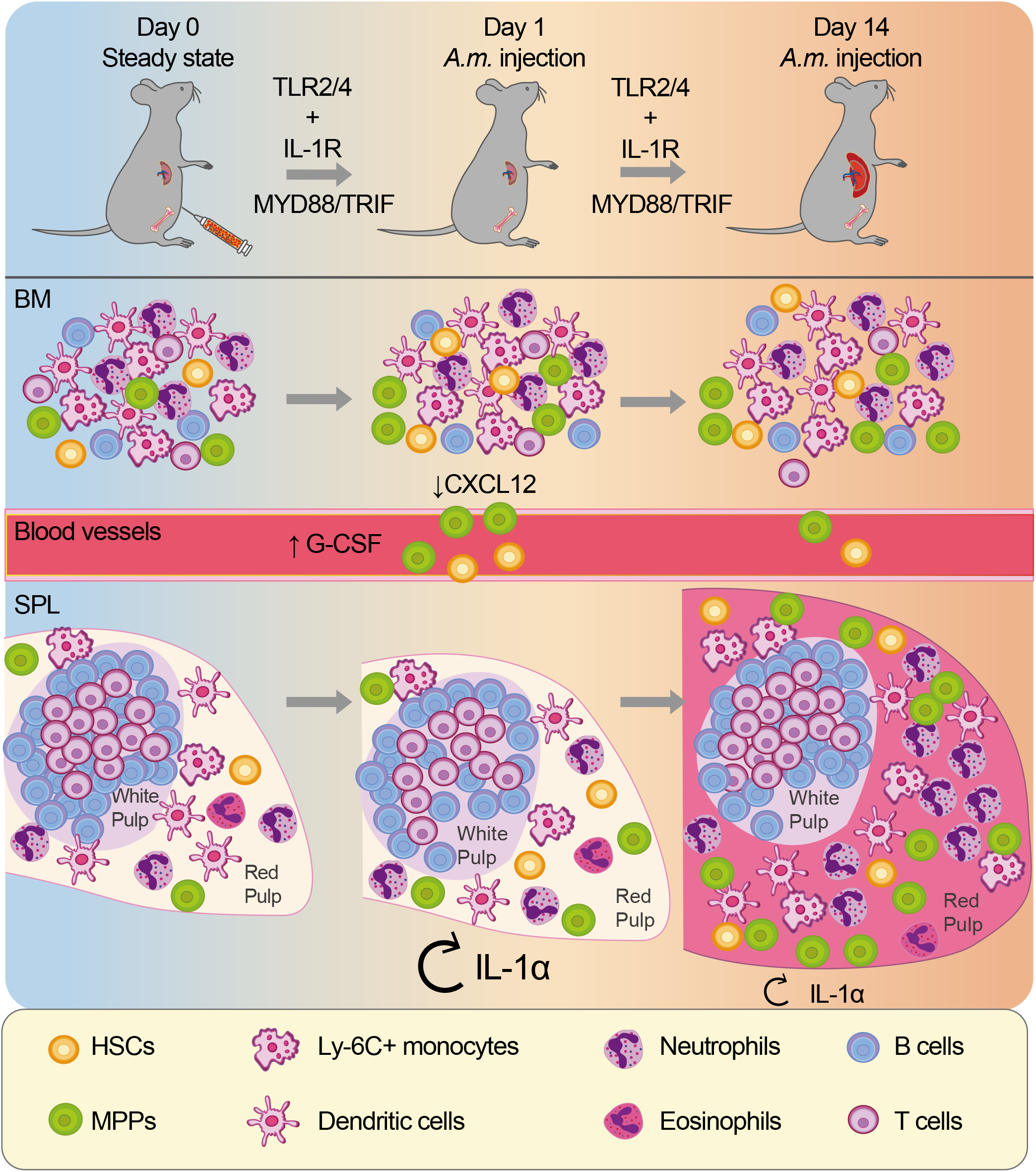
Extramedullary hematopoiesis is triggered by *A. m*. injection. The schematic model shows how the hematopoietic system responds to *A. m*. infection. At day 1 post *A. m*. injection, CXCL12 in the BM is downregulated, while the secretion of G-CSF in the PB increases, and mobilizes HSPCs from the BM to spleen. Concurrently, IL-1α secretion in the spleen is significantly elevated, and drives expansion of HSCs that have migrated to the spleen and their differentiation into myeloid cells. At day 14 post *A. m*. injection, persistent secretion of IL-1α and activation of TLR signals synergistically contribute to the constant expansion of functional HSPCs and their differentiation in spleen, ultimately leading to chronic EMH and splenomegaly.

Mechanistically, our data indicates chronic EMH to be dependent on downstream MYD88/TRIF signaling pathways, and partly mediated by TLR2/TLR4 and IL-1R (Fig. 5). Acute local section of IL-1α in the spleen appears largely dependent on TLR2/TLR4 signaling, but its chronic expression depends solely on MYD88/TRIF signaling pathways (Fig. 5A), implying that chronic IL-1α secretion is regulated by *A. m*. containing agents that stimulate only MYD88/TRIF-dependent innate immune signaling. This is supported by the fact that administration of either Pam3CSK4 or LPS alone was not sufficient to recapitulate chronic EMH (Fig. 4G-J). In contrast to previous reports stating that chronic LPS or IL-1*ε* stimulation damages HSC function^29, 37^, we did not see any obvious dysfunction in HSCs upon their transplantation (Fig. 3), also supporting IL-1α-dependent HSC expansion/differentiation. While the majority of splenic cell types that expand chronically depend on TLR2/TLR4 signaling, only those that chronically expand in the spleen depend on IL-1R signaling (Fig. 5G). What bacterial component in the *A. m*. lysate induces chronic EMH remains yet to be determined.

EMH occurs frequently during severe bacterial infection^18, 19^, metabolic stress^40^ and inflammatory bowel diseases (IBD)^30^, probably as a compensatory mechanism for ineffective BM hematopoiesis. Although EMH can be found in many organs, the spleen is considered to be one of the most common sites. This is likely because 1) the spleen plays a central role in host defense by combining both the innate and adaptive immune systems in an efficient way^41^ and 2) the spleen serves as a hematopoietic organ during embryonic development^42^. Similar to the BM niche, there is increasing evidence supporting spleen stromal cell subsets as key regulators of organ development and tissue regeneration^43^, albeit the underlying molecular mechanism is still unknown. Upon pathological EMH triggered by chemotherapy and G-CSF, SCF and CXCL12 secreted from splenic stromal cells are required for EMH^20^. Consistently, we found elevation of both SCF and CXCL12 in the spleen at day 1 post *A. m*. injection (data not shown), suggesting the splenic niche was disturbed by *A. m*.. Further study is necessary to address whether splenic niche cells secrets IL-1α to regulate *A. m*.-induced chronic EMH.

We found that late-onset chronic EMH in the spleen is mediated by innate immune signals, particularly TLR2/4 and IL-1R, with the latter possibly being regulated by local and persistent secretion of IL-1α upon *A. m*. stimulation. To our knowledge, IL-1α mediated EMH has not yet been reported. Abnormal activation of IL-1 family members is commonly found in certain chronic autoimmune-diseases such as rheumatoid arthritis (RA), atherosclerosis, and systemic lupus erythematosus (SLE)^44^. It has been recently shown that RA-derived HSCs are myeloid-biased via IL-1R-mediated signals^45^. It is also known that in human RA patients, aberrant hematopoiesis such as chronic anemia^46^, myeloid expansion as well as immuno-senescence^47^ are frequent causes for complications^48^. A subset of RA patients are associated with leukocytopenia and splenomegaly, known as Felty’s syndrome^49^.

Moreover, arthritis is frequently associated with IBD, a chronic inflammatory disorder highly related to intestinal barrier dysfunction and aberrant commensal microbiota distribution^50^. Given that *A. m*. is a mucin-degrading bacterium and *A. m*. can infiltrate into the circulation upon gut tissue damage, it can be readily speculated that *A. m*. might be involved in the pathogenesis of RA-associated EMH. Ours and other findings highlight that the cross-talk between multiple organs, or even horizontal interaction between different organisms, e.g., microbiome and host, may play a crucial role in the development of various immune-related diseases.

In conclusion, systemic stimulation with the membrane fraction of *A. m*., which is one of the abundant mucin-associated microbiota that is involved in various human pathologies induced late-onset EMH through innate immune signaling pathways mediated by TLR2/4 and IL-1R. These novel findings might be involved in the pathogenesis of IL-1-related autoimmune diseases and may lead to the development of novel therapies for their treatment.

## Acknowledgments

We would like to thank the International Core-facility of Advanced Life Science at Kumamoto University for their logistical and technical assistance and the Center for Animal Resources and Development at Kumamoto University for mice maintenance. We also thank Dr. Sayuri Nakata for her technical assistance.

## Author contribution

YW designed and performed the experiments, analyzed the data, and wrote the manuscript. TM and MS designed the experiments, and wrote the manuscript., GN and SF designed the experiments and prepared the bacterial lysates. YL and HT designed the experiments, supervised the research project and wrote the manuscript. All authors read and approved the final paper.

