## Supplemental method and figures for "*Akkermansia Muciniphila* induces chronic extramedullary hematopoiesis through cooperative IL-1R and TLR signals"

### Supplementary materials and methods

#### Mice

SPF C57BL/6 ($CD45.2^+$ background) and B6.SJL ($CD45.1^+$ background) mice were purchased from Japan SLC (Hamamatsu, Japan) and Jackson Laboratories (Bar Harbor, ME), respectively. CD45.1/$2^+$ mice were bred by crossing C57BL/6 ($CD45.2^+$) with B6.SJL ($CD45.1^+$). *Hlf*$^{tdTomato+}$ reporter mice ($CD45.1^+$) were kindly provided by Dr. Tomomasa Yokomizo (Kumamoto University)[39]. *Myd88;Trif*$^{-/-}$ and *TLR2;4*$^{-/-}$ mice ($CD45.2^+$) were purchased from Oriental BioService (Kyoto, Japan). All experiments were performed with 6-14 week-old mice. All mice were maintained at the Center for Animal Resources and Development at Kumamoto University. Experiments were approved by the Animal Care and Use Committee of Kumamoto University.

#### *A. muciniphila* preparation

*A. muciniphila* was cultured in GAM medium (#5422, Nissui Pharmaceutical, Tokyo, Japan) under anaerobic conditions. After 2 to 3 days of culture, bacteria were centrifuged at 15,000 x g for 5 minutes at 4°C and pellets was washed and resuspended in PBS. The bacterial suspension was then sonicated using a sonicator (Smurt, Microtec, Funabashi, Japan) with 30 second intervals on ice for at least 20 times. Efficiency was confirmed by gram-staining, and the bacterial suspension was ultracentrifuged at 100,000 x g for 60 minutes at 4°C (Optima XE-90, Beckman Coulter, Brea, CA) and resuspended in PBS. Protein concentration of the bacterial membrane fraction suspension was determined by Bio-Rad protein assay (#5000006, BioRad, Hercules, CA).

**Organ weight and blood counts**

Liver and spleen were isolated from treated mice and weighed. Peripheral blood was obtained from the orbital vein using heparin-coated capillaries (HIRSCHMANN, Eberstadt, Germany). Hematological parameters were assessed using a hematology analyzer (Celltac α MEK-6358, Nihon Kohden, Tokyo, Japan).

**Treatment**

Mice were injected with the following reagents intraperitoneally (*i.p.*): 200μg of LPS, corresponding to 200EU LPS-EB Ultrapure (#14954-81, Invivogen, San Diego, CA), 200μg of Pam3CSK4 (#14954-61, Invivogen), 200μg of *A.m.* or PBS as a vehicle control. For the IL-1R inhibition experiment, mice were *i.p.* injected, daily with 37μg of Anakinra (Kineret, IL-1ra, Swedish Orphan Biovitrum AB, Stockholm, Sweden) over 2 weeks until the mice were sacrificed for analysis. *A.m.* was injected once after the first Anakinra injection in this experiment.

**FACS analysis**

BM and spleen were harvested and incubated with the following biotinylated antibodies against the lineage (Lin) markers: B220 (RA3-6B2), CD3ε (145-2C11), CD4 (GK1.5), CD8α (53-6.7), NK1.1 (PK136), CD11b (M1/70), Ter119 (Ter119), and Gr-1 (RB6-8C5) together with the fluorescence-conjugated antibodies: c-Kit (2B8), Sca-1 (D7), CD34 (HM34), Flt3 (A2F10), CD150 (TC15-12F12.2), CD48 (HM48-1), CD16/32 (93), IL-7Rα (A7R34) and CD86(GL-1), and for mature cell analysis with: B220 (RA3-6B2), CD3ε (145-2C11), F4/80 (BM8), Ly6G (1A8), Ly6C (HK1.4) and CD11b (M1/70). Cells were analyzed on FACS Canto II or FACS Aria III flow cytometers (BD

Biosciences, Franklin Lakes, NJ). Data was analyzed using FlowJo (BD Biosciences).

**Competitive repopulation assays**

$3x10^5$ of whole BM (WBM) cells and $1x10^6$ of whole splenocytes were isolated from either *A.m.*- or PBS-treated WT mice ($CD45.1^+$) at 14 days post injection. These cells were mixed with $3x10^5$ WBM competitor cells isolated from WT mice (CD45.1/CD45.2) and transplanted intravenously into lethally-irradiated (10Gy) 8-10 weeks old WT mice ($CD45.2^+$). Donor chimerism in PB was assessed every 4 weeks up to 16 weeks post transplantation. Mice were sacrificed at 20 weeks after transplantation and donor chimerism was assessed in the BM.

**Measurement of inflammation- or HSC niche-related factors**

For serum collection, PB was obtained via the orbital vein using glass capillaries without anti-coagulants and left at room temperature for 20 minutes followed by centrifugation at 1,000 x *g* for 15 minutes. To obtain BM fluid, two tibiae were flushed out with 0.5ml BSA/PBS (0.1%), followed by centrifugation at 3,500 rpm for 5 minutes at 4°C. For spleen fluid, a piece of spleen (5 mg) was mashed between a pair of glass slides in 0.1ml BSA/PBS (0.1%), followed by centrifugation at 3,500 rpm for 5 minutes at 4°C. The splenocyte lysate was obtained by lysing cells with the same volume of 500μM 1x RIPA buffer (Abcam, Cambridge, UK). Samples were all stored at -80 C° until cytokine measurement after collection. G-CSF was quantified by Mouse G-CSF ELISA Kit (#KE10025, Proteintech, Rosemont, IL) according to the manufacturer's instructions. Other cytokines were measured with Legendplex™ Mouse Inflammation Panel (#740150, BioLegend) and Legendplex™ Mouse HSC Panel (#740150, BioLegend) according to the manufacturer's instructions.

**Spleen imaging**

Spleen samples harvested from Hlf$^{tdTomato+}$ reporter mice at 14 days post *A.m.* lysate injection were fixed in 4% PFA at 4°C overnight, washed in PBS for 30 minutes and cryoprotected with 30% sucrose in PBS at 4°C overnight. Samples were imbedded in O.C.T compound (Sakura Finetek Japan, Tokyo, Japan) and frozen, followed by sectioning (20µm) using a Leica CM1950 cryostat (Leica Biosystems, Wezlar, Germany). Sectioned samples were washed with PBS 3 times to remove excess O.C.T compound and incubated with the following primary antibodies at 4°C overnight: RFP (200-301-379, ROCKLAND, Limerick, PA), c-Kit (AF1356, R&D systems, Minneapolis, MN), Gr-1 (RB6-8C5, 16-5931-81, Thermo Fisher Scientific). Samples were washed with PBS 3 times and stained with the following secondary antibodies at room temperature for 1 hour: Donkey anti-Rabbit IgG (H+L) conjugated with Cy-3 and donkey anti-Rat IgG (H+L) conjugated with Alexa Fluor 647 (Jackson ImmunoResearch, West Grove, PA). directly fluorescent conjugated antibodies were used for F4/80-PE (BM8, Thermo Fisher Scientific) and Ly6G-AF647 (1A8, BioLegend) staining. Images were obtained with a Leica CS SP8 DLS confocal microscope (Leica Biosystems). Three-dimensional (3D) reconstructions were generated from z-stack images using the microscopy imaging analysis software Imaris (Bitplane, UK).

**Supplemental Figure Legends**

**Figure S1. FACS analysis of HSPCs in the BM and spleen of *A.m.*-treated mice**

(A) Gating strategy for HSPC subpopulation analysis. All events were gated for FSC-A against SSC-A. Singlets were selected using FSC-H against FSC-W and SSC-H against SSC-W. Lineage-committed cells were excluded from live cells. Within the lineage-negative ($Lin^-$) population, IL-$7R\alpha^+$ and $c\text{-}Kit^{mid}$ cells were defined as CLPs. The LK population was gated as single positive for c-Kit and negative for Sca-1, while LSK cells were positive for both c-Kit and Sca-1. LSK ($Lin^-$ $Sca^-1^+Kit^+$) cells were further divided into $HSC^{LT}$ ($CD150^+CD48^-$ LSK), $HSC^{ST}$ ($CD150^-CD48^-$ LSK), MPP2 ($CD150^+CD48^+$LSK) and MPP3/4 ($CD150^+CD48^+$LSK). MPP3/4 were characterized as $CD135^-$ MPP3 or $CD135^+$ MPP4. LK ($Lin^-Sca\text{-}1^-Kit^+$) cells were divided into MEP ($CD34^-CD16/32^{low}$LK), GMP ($CD34^+CD16/32^+$LK), and CMP ($CD34^+CD16/32^+$LK). (B-C) Representative FACS plots (B) and frequency (C) of mature cell fractions in BM at 14 days after PBS or *A.m.* injection into WT mice (n=12-13 from 7 independent experiments). (D-E) Representative FACS plots (D) and frequency (E) of LK, LSK and CLP in $Lin^-$ BM at 14 days after PBS or *A.m.* injection into WT mice (n=5-8 from 4 independent experiments). (F-G) Representative images of spleen from PBS- or *A.m.*-treated mice at day 14. (F) c-Kit, green; Gr-1, yellow; DAPI, blue, (G) F4/80, red; Ly-6G, yellow; DAPI, blue. Scale bar indicates 100μm. (H) Percentage of HSPC subpopulation in $Lin^-$ cells of spleen and BM at day 14 after *A.m.* injection via *i.p.* or *i.v.* routes into WT mice (n=3 from 2 independent experiments).

**Figure S2. Time-course kinetic analysis revealed two distinct waves of HSPC increase in the spleen of *A.m.*-injected mice**

(A) Absolute number of mature cell fractions in the spleen of PBS- or *A.m.*-injected WT mice. (B) Representative FACS plots of HSPC subfractions in $Lin^-$ cells from the BM (upper), spleen (middle), and PB (lower) at indicated days after PBS or *A.m.* injection into WT mice. (C) Absolute number of HSPC subfractions in the BM of PBS- or *A.m.*-injected WT mice (3-14 mice from 3-8 independent experiments). (D-E) Representative FACS plots (D) and frequency (E) of CD86- or Sca-1-defined HSPC in $Lin^-$ BM at day 1 and 14 after PBS or *A.m.* injection into WT mice (n=3-8 from 5 independent experiments).

**Figure S3.** ***A.m.*** **induced HSPC accumulation in the BM through innate immune signals**

(A) Representative FACS plots of $Lin^-$ BM cells from PBS- or *A.m.*-treated WT, *Tlr2/4* or *Myd88/Trif* DKO mice at day 14 post injection. (B) Absolute number of HSPC subpopulations in BM WT mice injected with PBS (WT-PBS, black box) or *A.m.* (WT-*A.m.*, red in black box), *Tlr2;4* DKO mice injected with PBS (*Tlr2;4* DKO-PBS, blue box) or *A.m.* (*Tlr2;4* DKO-*A.m.*, red in blue box), or *Myd88/Trif* DKO mice injected with PBS (*Myd88;Trif* DKO-PBS, green box) or *A.m.* (*Myd88;Trif* DKO-*A.m.*, red in green box) at day 14 post injection (n=4-11 from 4 independent experiments). (C) Representative FACS plots of $Lin^-$ cells in the BM at day 14 after PBS, 200μg *A.m.*, 200μg LPS or 200μg Pam3CSK4 injection into WT mice. (D-E) Absolute cell numbers of whole BM (D) and HSPC subpopulations (E) in the BM at day 14 after PBS, *A.m.*, LPS or Pam3CSK4 injection into WT mice. (n=4-7 from 2 independent experiments).

**Figure S4. Local secretion of IL-1α contributed to the chronic expansion of HSPCs in the spleen**

(A) Concentration of IL-1α, IL-1β, TNF-α and IFN-γ in spleen cell lysate isolated from WT, *Tlr2;4* DKO and *Myd88;Trif* DKO mice treated with PBS or *A.m.* at day 1 and 14 post injection (n=3-6 from 3 independent experiments). (B) Concentration of inflammation-related (upper) and HSC niche-related (lower) factors in the serum of WT mice treated with PBS or *A.m.* at day 1 and 14 post injection (n=6-7 from 4 independent experiments). (C-D) Concentration of HSC niche-related factors in spleen cell lysate (C) and spleen fluid (D) isolated from WT mice treated with PBS or *A.m.* at day 14 post injection (n=6-7 from 4 independent experiments). (E-F) Representative FACS plots (E) and absolute cell number (F) of HSPC subpopulations in the BM at day 14 post injection (n=5-6 from 4 independent experiments). Dashed lines indicate absolute cell numbers for each HSPC fraction in PBS-treated BM.

Wang et al. Figure S1 FACS analysis of HSPCs in BM and spleen of *A.m.*-treated mice.

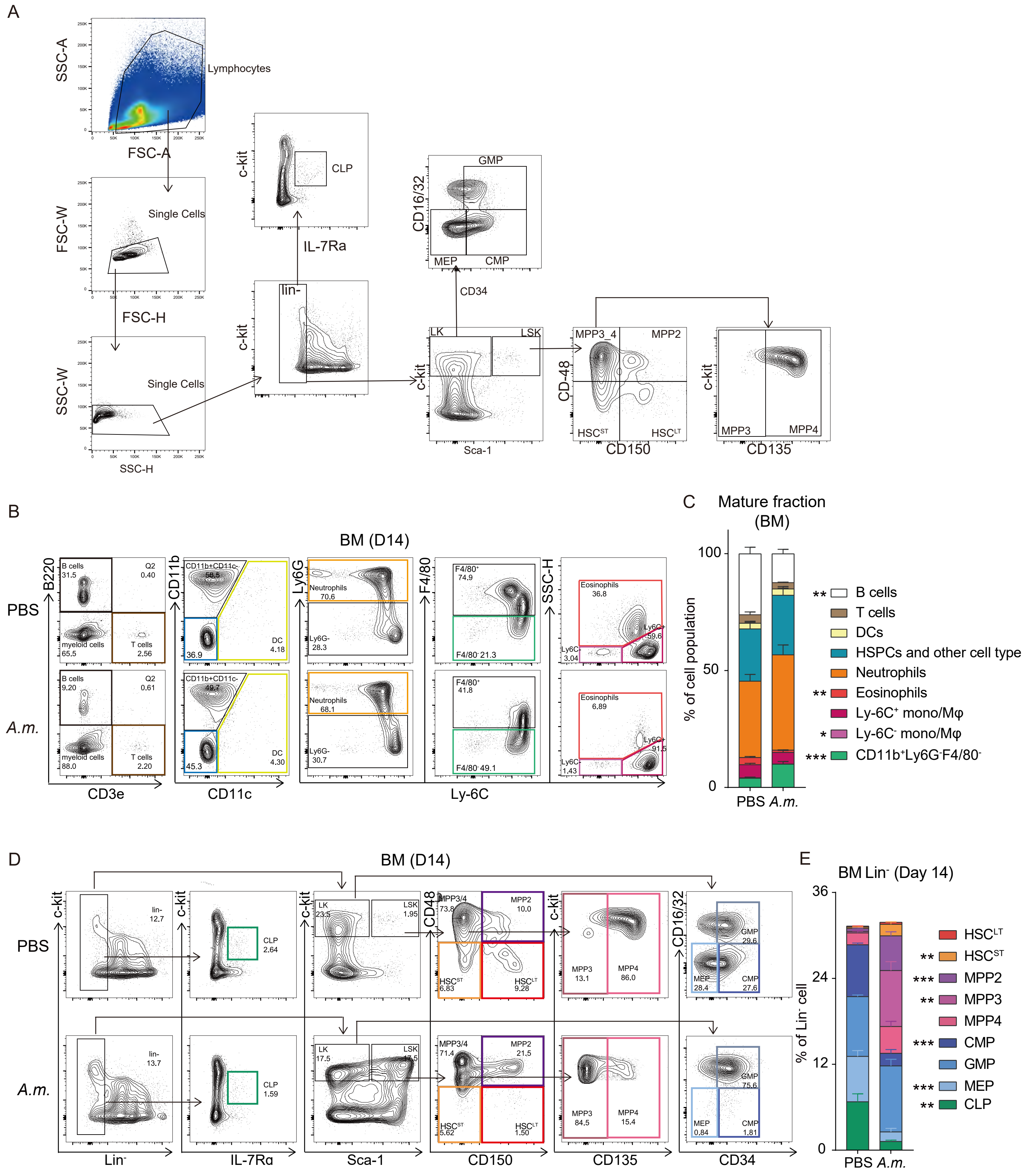

Wang et al. Figure S1 FACS analysis of HSPCs in BM and spleen of *A.m.*-treated mice. (Continued)

F

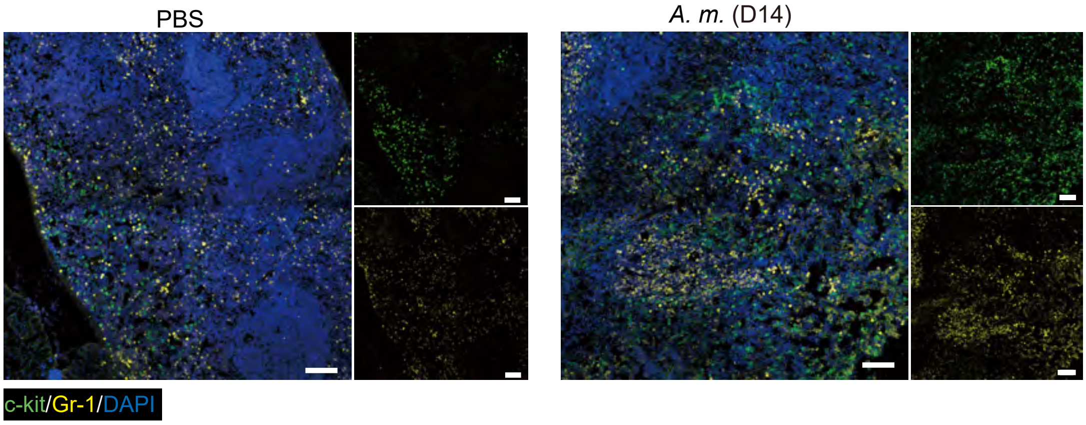


G

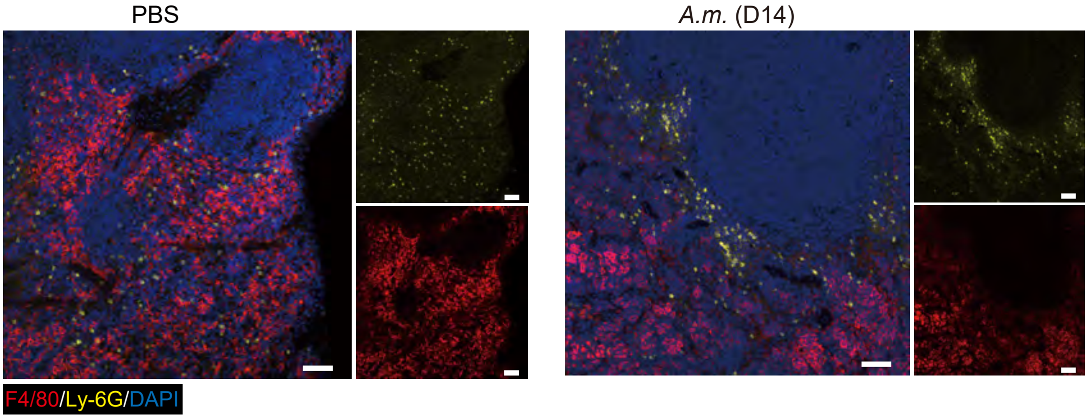


H

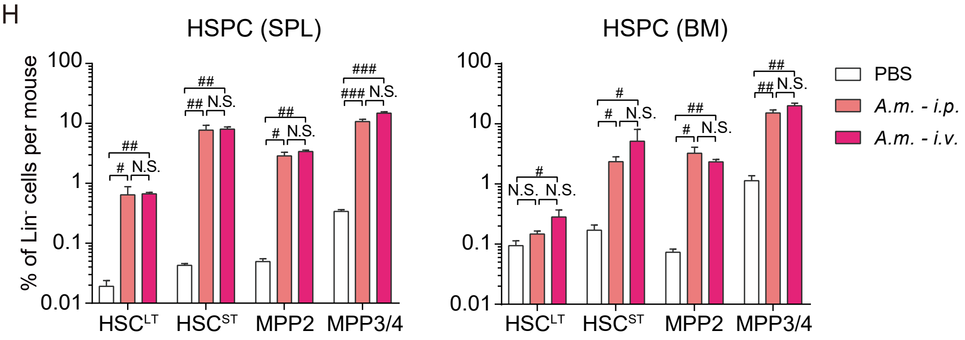

Wang et al. Figure S2 Time-course kinetic analysis revealed two differential HSPC waves in *A. m.*-injected spleen.

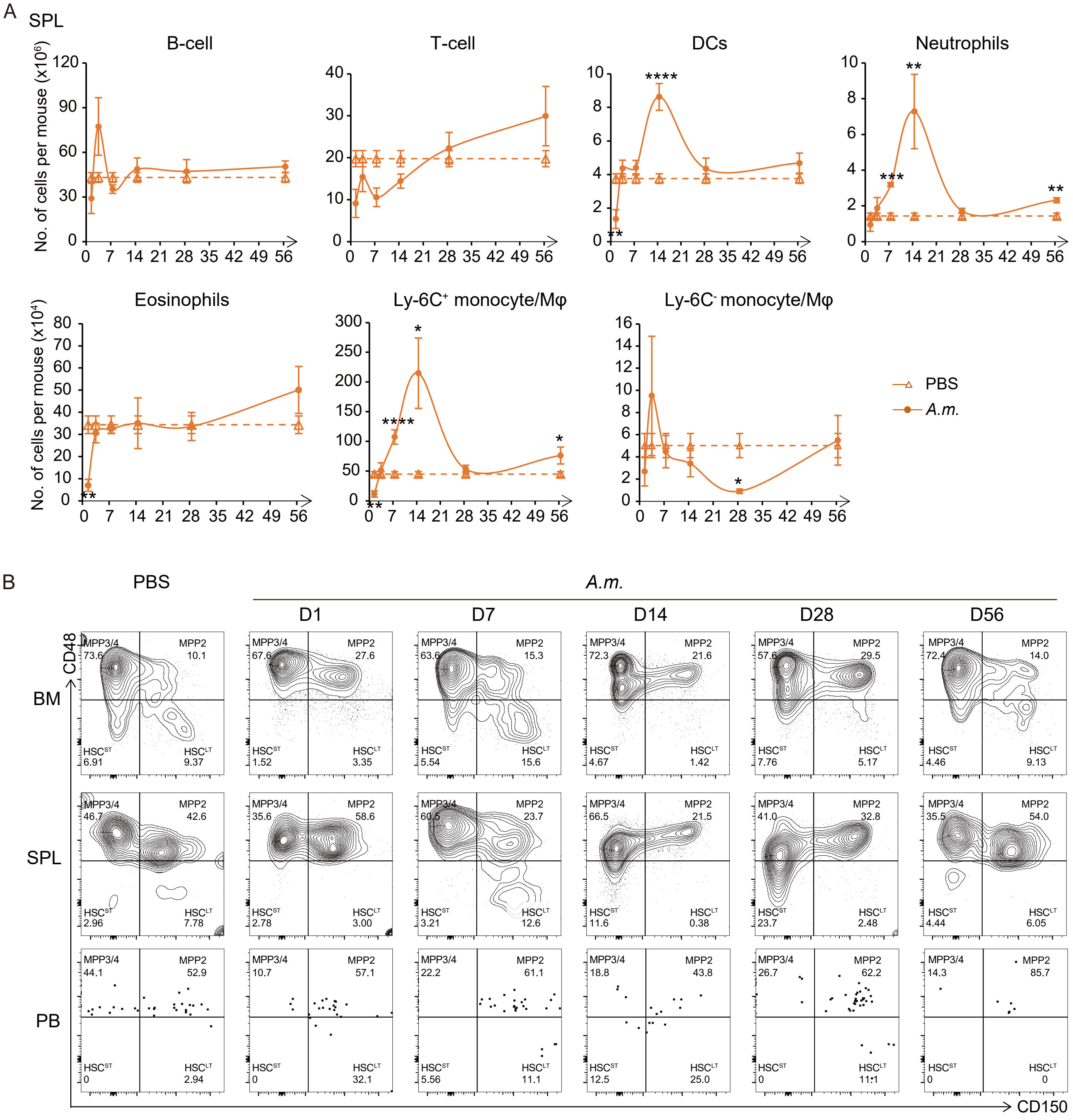

Wang et al. Figure S2 Time-course kinetic analysis revealed two differential HSPC waves in *A. m.*-injected spleen. (Continued)

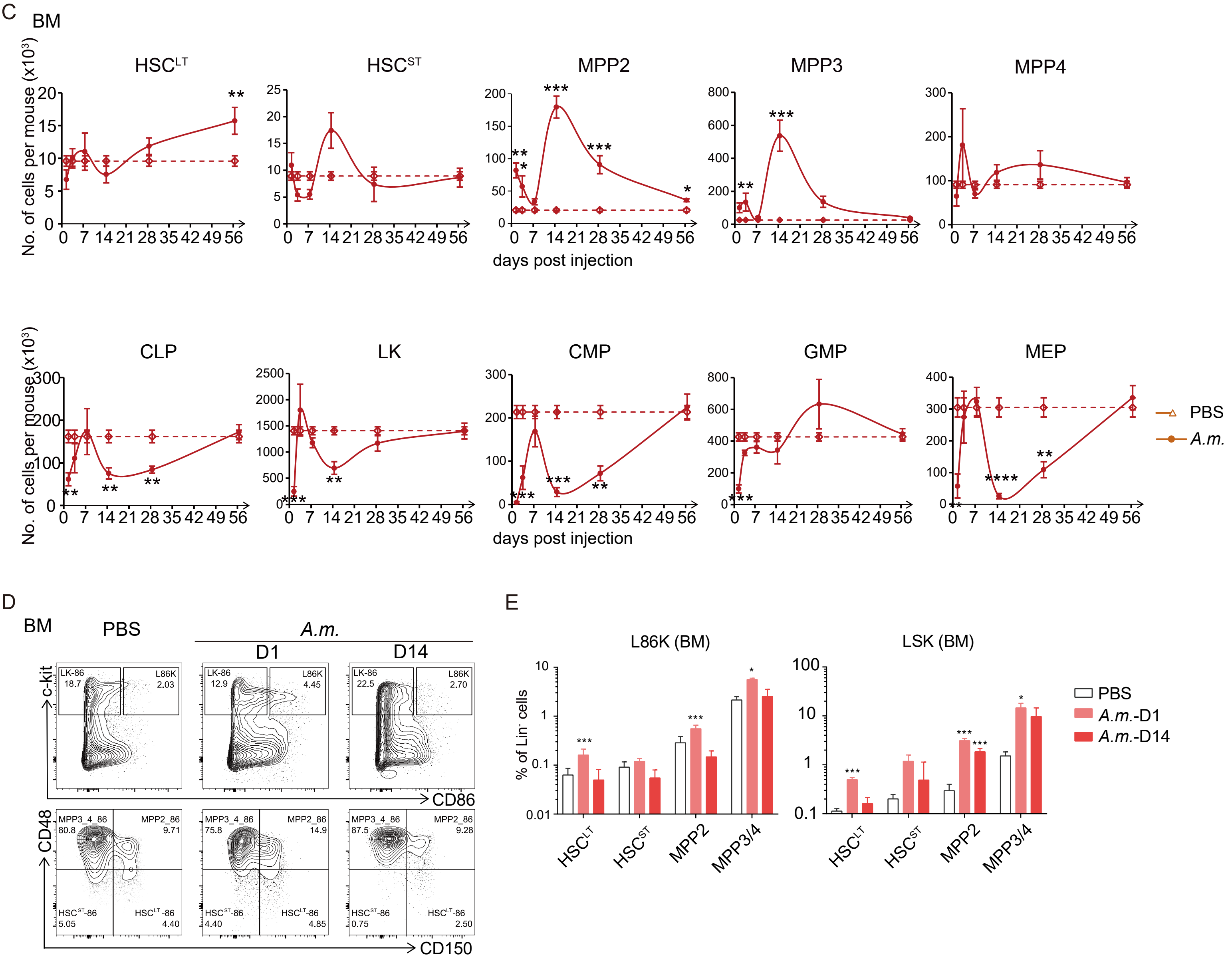

Wang et al. Figure S3 *A. m.* induced HSPC expansion in the spleen is psrtially mediated by TLR2/4 signals and entirely by MYD88/TRIF signals.

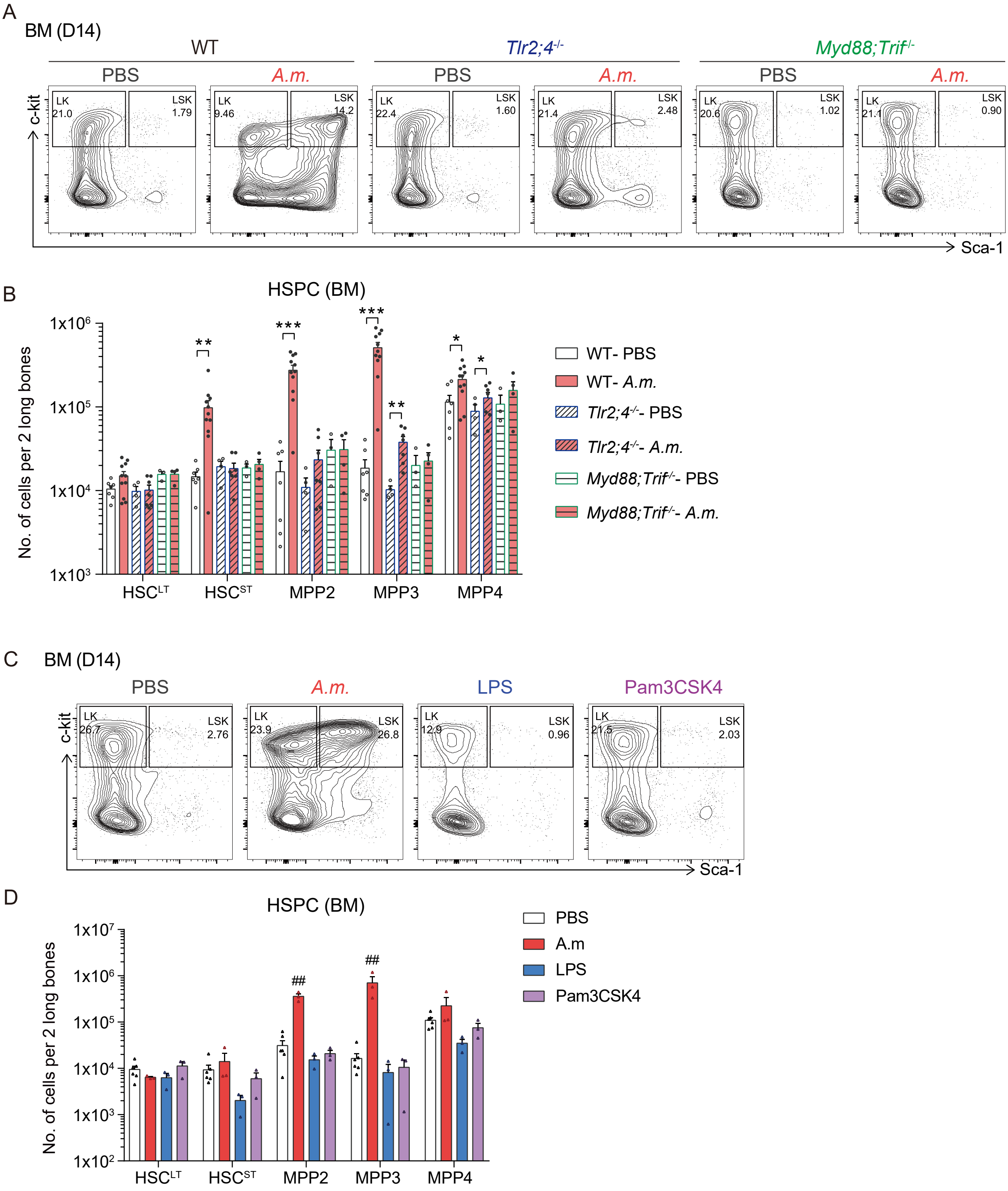

Wang et al. Figure S4 Sustained elevation of splenic IL-1α production contributes to TLR2/4-independent chronic EMH.

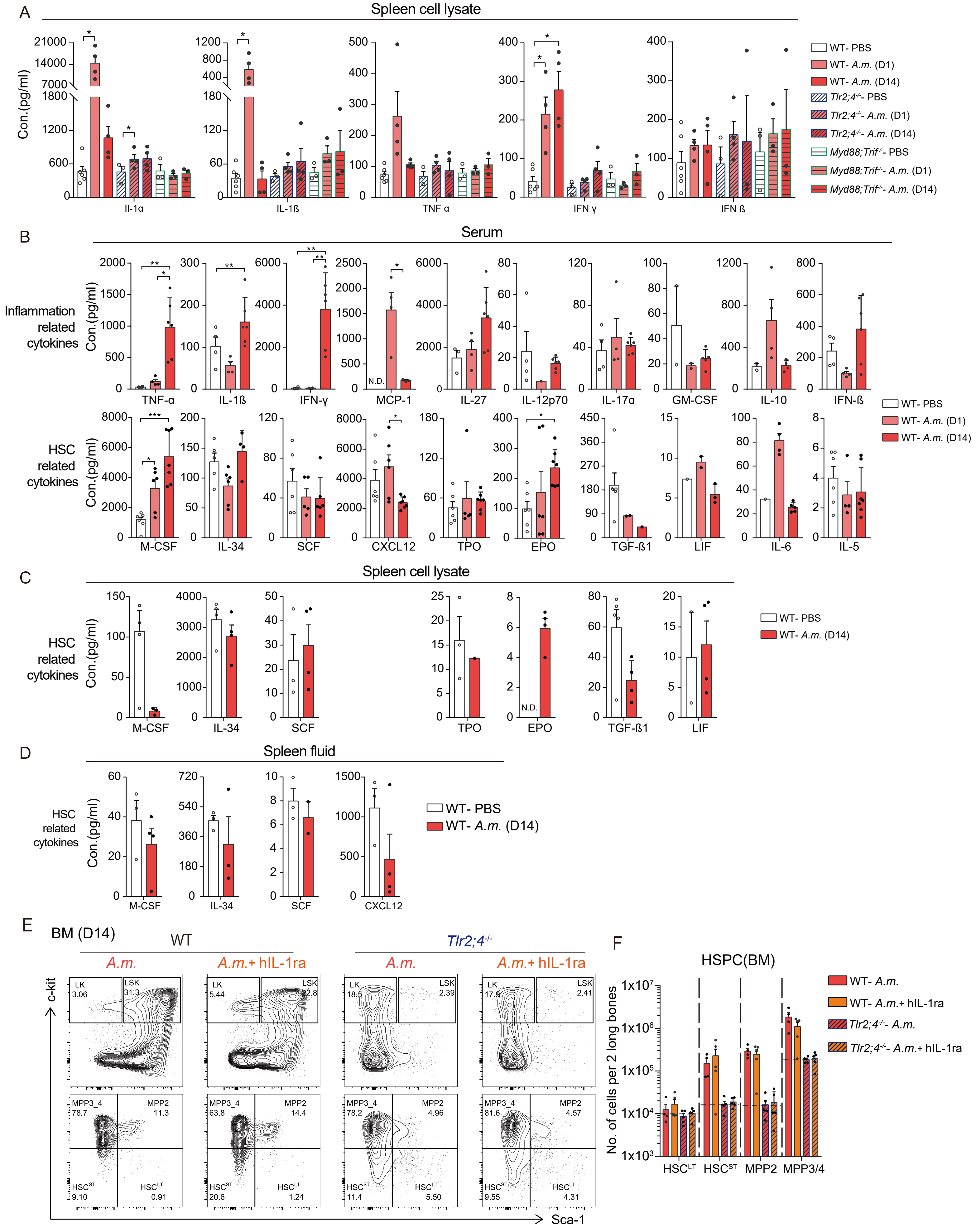
